## Supplemental information for "*Ex vivo* Expansion Potential of Murine Hematopoietic Stem Cells: A Rare Property Only Partially Predicted by Phenotype"

### SUPPLEMENTAL TABLES

#### Supplemental Table 1. Contents of murine HSC media for *in vitro* culture

| | Final concentration | Provider |
|---|---|---|
| Insulin-transferrin-selenium-ethanolamine (ITSX) | 1 x | Gibco |
| Penicillin/streptomycin/Glutamine (P/S/G) | 1 x | Gibco |
| HEPES | 10 mM | Gibco |
| Polyvinyl alcohol (PVA) | 1 mg/ml | Sigma |
| Animal free stem-cell factor (SCF) | 10 ng/ml | Peprotech |
| Animal free thrombopoietin (TPO) | 100 ng/ml | Peprotech |
| In Ham's F-12 Nutrient Mix (Gibco) | | |

#### Supplemental Table 2. Lineage cocktail (Biotinylated)

| Surface marker | Coupled dye | Clone | Dilution | Provider |
|---|---|---|---|---|
| B220 | Biotin | RA3-6B2 | 1:200 | Sony |
| CD3 | Biotin | 17A2 | 1:100 | Sony |
| Gr-1 | Biotin | RB6-8C5 | 1:400 | Sony |
| Nk1.1 | Biotin | PK136 | 1:400 | BD |
| Ter119 | Biotin | TER-119 | 1:400 | Sony |

#### Supplemental Table 3. Lineage cocktail (PE-Cy5)

| Surface marker | Coupled dye | Clone | Dilution | Provider |
|---|---|---|---|---|
| B220 | PE-Cy5 | RA3-6B2 | 1:200 | Sony |
| CD3 | PE-Cy5 | 145-2C11 | 1:400 | Sony |
| Gr-1 | PE-Cy5 | RB6-8C5 | 1:400 | Sony |
| Nk1.1 | PE-Cy5 | PK136 | 1:400 | Sony |
| Ter119 | PE-Cy5 | TER-119 | 1:400 | Sony |

**Supplemental Table 4. Antibody mixture for BM or FL cHSC sorting**

| Surface marker | Coupled dye | Clone | Dilution | Provider |
|---|---|---|---|---|
| Lineage cocktail | As outlined in Table 2. | | | |
| CD48 | FITC | HM48-1 | 1:200 | Sony |
| CD150 | PE-Cy7 | TC15-12F12.2 | 1:200 | Sony |
| CD201 | PE | RCR-16 | 1:200 | Sony |
| Sca1 | PB | E13-161.7 | 1:200 | BioLegend |

* CD41-PerCP-eFluor710 (1:100, Clone eBioMWReg30, eBioscience Cat. #46-0411-82) was included if mentioned in the results.

**Supplemental Table 5. Antibody mixture for cHSC culture analysis and sorting**

| Surface marker | Coupled dye | Clone | Dilution | Provider |
|---|---|---|---|---|
| Lineage cocktail | As outlined in Table 3. | | | |
| CD48 | AF700 | HM48-1 | 1:100 | Sony |
| CD150 | PE-Cy7 | TC15-12F12.2 | 1:200 | Sony |
| CD201 | PE | RCR-16 | 1:200 | Sony |
| cKit | APCeFluor780 | 2B8 | 1:100 | eBioscience |
| Fcer1a | FITC | MAR-1 | 1:200 | eBioscience |
| Sca1 | PB | E13-161.7 | 1:200 | BioLegend |

* In case of analysis of expansion from Fgd5-ZsGreen labeled HSCs, Fcer1a-FITC was excluded.

**Supplemental Table 6. Antibody mixture for PB chimerism analysis**

| Surface marker | Coupled dye | Clone | Dilution | Provider |
|---|---|---|---|---|
| CD3 | AF700 | 17A2 | 1:200 | Sony |
| CD11b | APC | M1/70 | 1:200 | Sony |
| CD19 | PE-Cy7 | 6D5 | 1:400 | Sony |
| CD45.1 | BV650 | A20 | 1:100 | Sony |
| CD45.2 | BV785 | 104 | 1:100 | Sony |
| Gr-1 | PE | RB6-8C5 | 1:400 | Sony |
| Nk1.1 | PB | PK136 | 1:200 | Sony |
| Ter119 | PerCP-Cy5.5 | TER-119 | 1:200 | Sony |

**Supplemental Table 7. Antibody mixture for BM cHSC chimerism analysis**

| Surface marker | Coupled dye | Clone | Dilution | Provider |
|---|---|---|---|---|
| Lineage cocktail | As outlined in Table 3. | | | |
| CD45.1 | BV650 | A20 | 1:100 | Sony |
| CD45.2 | BV785 | 104 | 1:100 | Sony |
| CD48 | AF700 | HM48-1 | 1:100 | Sony |
| CD150 | PE-Cy7 | TC15-12F12.2 | 1:200 | Sony |
| CD201 | PE | RCR-16 | 1:200 | Sony |
| cKit | APCeFluor780 | 2B8 | 1:100 | eBioscience |
| Sca1 | PB | E13-161.7 | 1:200 | BioLegend |

**Supplemental Table 8. Antibody mixture for culture CTV analysis**

| Surface marker | Coupled dye | Clone | Dilution | Provider |
|---|---|---|---|---|
| Lineage cocktail | As outlined in Table 3. | | | |
| CD48 | APC | HM48-1 | 1:200 | Sony |
| CD150 | PE-Cy7 | TC15-12F12.2 | 1:200 | Sony |
| CD201 | PE | RCR-16 | 1:200 | Sony |
| cKit | APCeFluor780 | 2B8 | 1:100 | eBioscience |
| Fcer1a | FITC | MAR-1 | 1:200 | eBioscience |
| Sca1 | BV711 | D7 | 1:200 | Sony |

**Supplemental Table 9. Antibody mixture for BM CTV analysis**

| Surface marker | Coupled dye | Clone | Dilution | Provider |
|---|---|---|---|---|
| Lineage cocktail | As outlined in Table 3. | | | |
| CD45.1 | BV650 | A20 | 1:100 | Sony |
| CD45.2 | BV785 | 104 | 1:100 | Sony |
| CD48 | FITC | HM48-1 | 1:200 | Sony |
| CD135 | PE | A2F10 | 1:100 | Sony |
| CD150 | PE-Cy7 | TC15-12F12.2 | 1:200 | Sony |
| CD201 | APC | eBio1560 | 1:200 | eBioscience |
| Sca1 | BV711 | D7 | 1:200 | Sony |

**Supplemental Table 10. Antibody mixture for Spleen CTV analysis**

| Surface marker | Coupled dye | Clone | Dilution | Provider |
|---|---|---|---|---|
| CD4 | APC-Cy7 | GK1.5 | 1:200 | Sony |
| CD11b | FITC | M1/70 | 1:400 | BD |
| CD19 | PE-Cy7 | 6D5 | 1:400 | Sony |
| CD45.1 | BV650 | A20 | 1:100 | Sony |
| CD45.2 | BV785 | 104 | 1:100 | Sony |
| Ter119 | PE-Cy5 | TER-119 | 1:400 | Sony |

**Supplemental Table 10. List of primers**

| Primer pair | Sequence (5' – 3') |
|---|---|
| Pre-culture Forward | TCGTCGGCAGCGTCAGATGTGTATAAGAGACAGGAAGCTGC GCCTGTCATC |
| Pre-culture Reverse | GTCTCGTGGGCTCGGAGATGTGTATAAGAGACAGGTGAACC GCATCGAGCTG |
| Post-culture Forward | TCGTCGGCAGCGTCAGATGTGTATAAGAGACAGTGGAGAAC CACCTTGTTGG |
| Post-culture Reverse | GTCTCGTGGGCTCGGAGATGTGTATAAGAGACAGTGCATGG CGGTAATACGGT |
| P5 index N101 | AATGATACGGCGACCACCGAGATCTACAC **TAGATCGC** TCGTCGGCAGCGTC |
| P5 index S502 | AATGATACGGCGACCACCGAGATCTACAC **CTCTCTAT** TCGTCGGCAGCGTC |
| P5 index S503 | AATGATACGGCGACCACCGAGATCTACAC **TATCCTCT** TCGTCGGCAGCGTC |
| P7 index N701 | CAAGCAGAAGACGGCATACGAGAT **TCGCCTTA** GTCTCGTGGGCTCGG |
| P7 index N901 | CAAGCAGAAGACGGCATACGAGAT **AACGTGAT** GTCTCGTGGGCTCGG |
| P7 index N902 | CAAGCAGAAGACGGCATACGAGAT **AAACATCG** GTCTCGTGGGCTCGG |
| P7 index N903 | CAAGCAGAAGACGGCATACGAGAT **ATGCCTAA** GTCTCGTGGGCTCGG |
| P7 index N904 | CAAGCAGAAGACGGCATACGAGAT **AGTGGTCA** GTCTCGTGGGCTCGG |

### SUPPLEMENTAL FIGURES

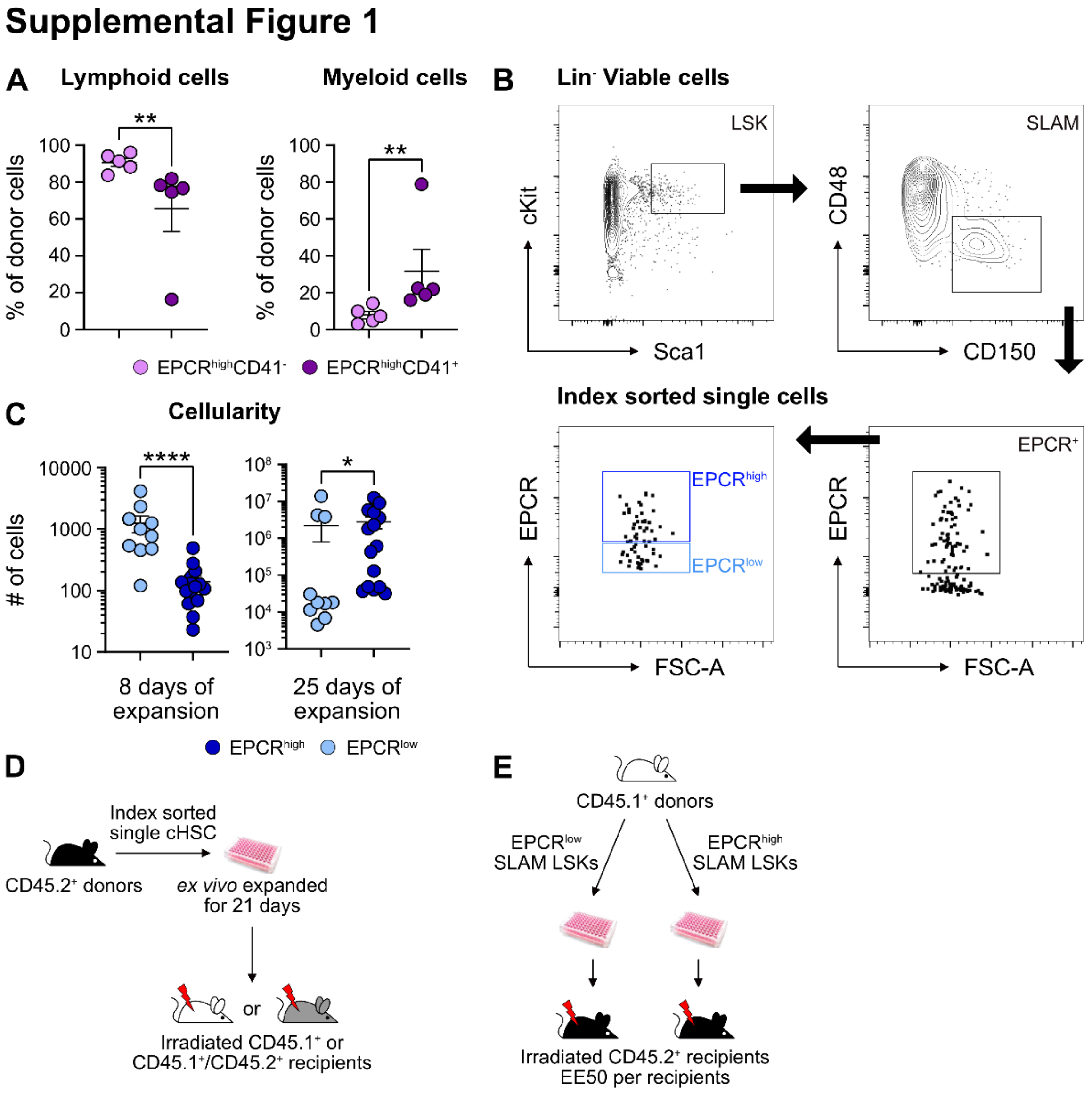


**Supplemental Figure 1. The PB lineage output from EPCR$^{high}$CD41$^{-}$ and EPCR$^{high}$CD41$^{+}$ cells after transplantation.**

(A) The distribution of lymphoid (pooled B and T cells) and myeloid cells out of the test-cell derived reconstitution 16 weeks post-transplantation.

(B) Index sorting strategy for single cell cultures.

(C) Overall cell expansion from one index sorted SLAM LSKs after 8 or 25 days of *ex vivo* culture. Cultures were separated into two groups based on EPCR expression level.

(D) Strategy to assess repopulating and radioprotection ability of *ex vivo* expanded cells from index sorted single SLAM LSKs.

(E) Strategy to assess repopulating and radioprotection ability of cells *ex vivo* expanded from 50 $EPCR^{high}$ or $EPCR^{low}$ SLAM LSKs.

Data points depict values in individual recipients or individual cultures. Error bars denote SEM. The asterisks indicate significant differences. *, $p < 0.05$; **, $p < 0.01$; ****, $p < 0.0001$.

### Supplemental Figure 2

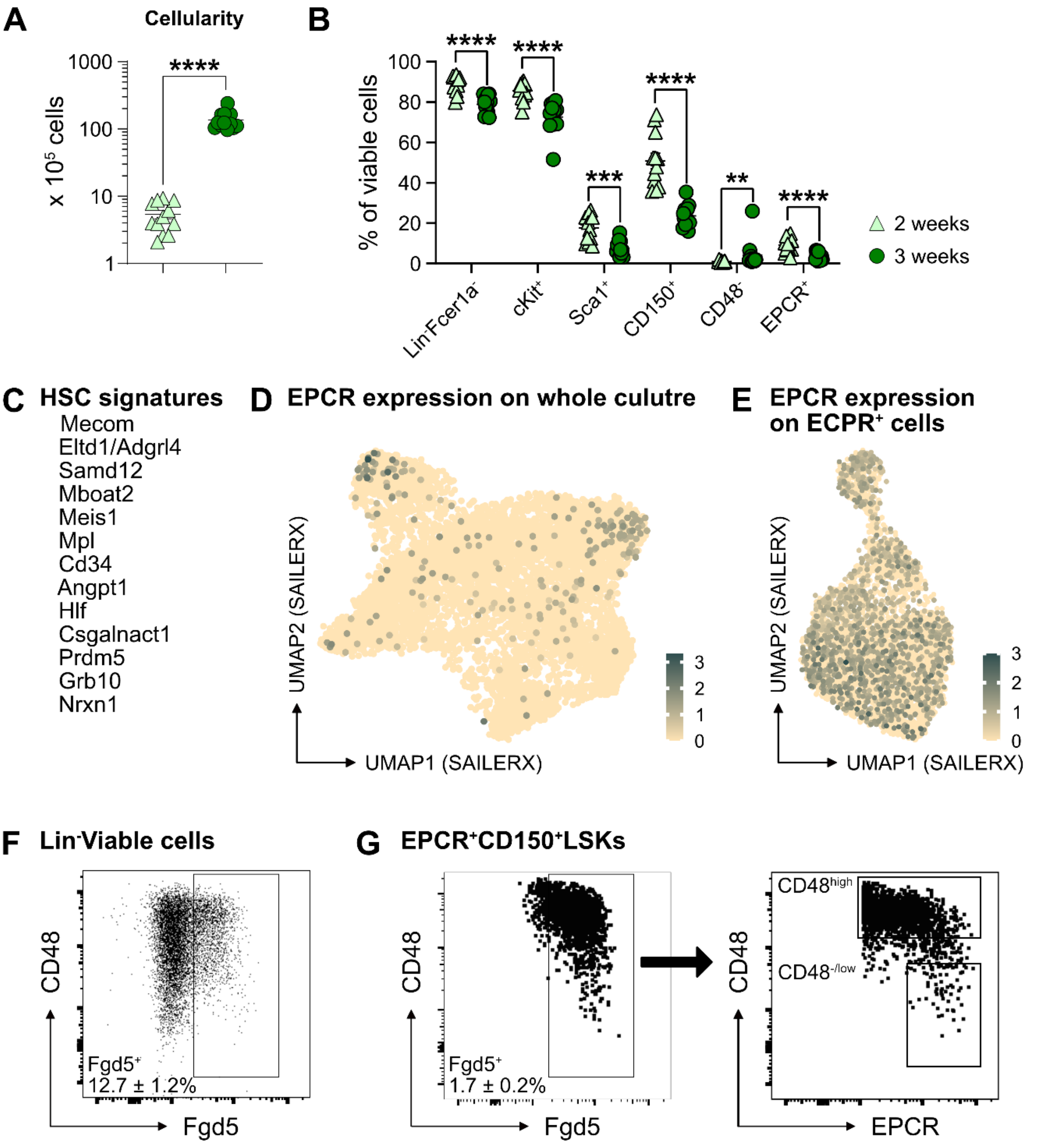


**Supplemental Figure 2. Heterogeneity of *ex vivo* expanded cHSCs.**

(A) Overall cell expansion from 50 EPCR$^{high}$ SLAM LSKs after 14- or 21-days of *ex vivo* culture.

(B) Frequency of cells expressing different stem cell markers in *ex vivo* cultures following 14 or 21 days of culture. Data points depict values from individual cultures initiated from 50 cHSCs.

(C) List of signatures used to define cHSCs.

(D) and (E) Expression of EPCR on whole culture and EPCR$^+$ cells.

(F) and (G) Representative FACS plots of cells expanded 14 or 21 days in *ex vivo* cultures using Fgd5-ZsGreen reporter cells, respectively. Mean values demonstrate the frequency of Fgd5$^+$Lin$^-$Fcer1a$^-$ or Fgd5$^+$EPCR$^+$CD150$^+$Fcer1a$^-$LSK cells, respectively. Mean ± SEM value was calculated from 15 individual cultures initiated from 50 cHSCs.

Error bars denote SEM. The asterisks indicate significant differences. **, $p < 0.01$; ***, $p < 0.001$; ****, $p < 0.0001$.

### Supplemental Figure 3

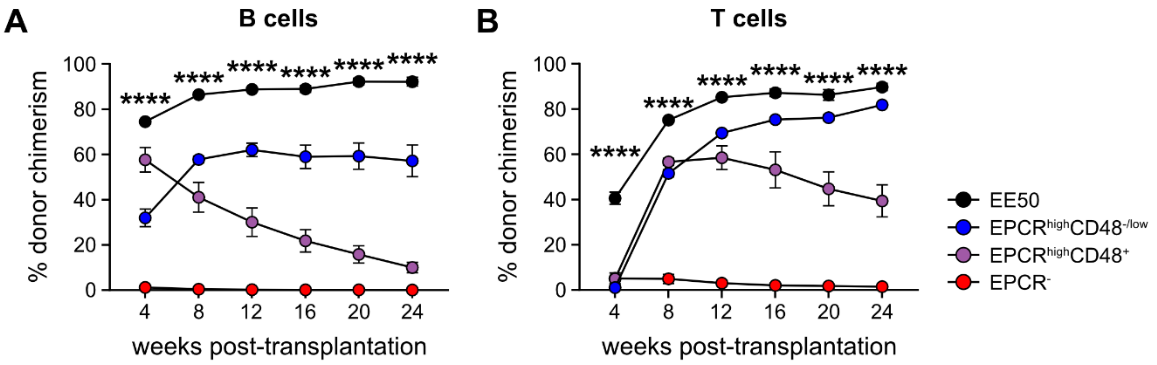


**Supplemental Figure 3. B and T cell chimerism in PB after transplantation.**

(A) Test-cell derived chimerism in PB B cells over 24 weeks post-transplantation.

(B) Test-cell derived chimerism in PB T cells over 24 weeks post-transplantation.

Data represent mean values (n = 5 per group). Error bars denote SEM. A one-way ANOVA test was applied and the asterisks indicate significant differences among the four groups. ****, $p < 0.0001$.

### Supplemental Figure 4

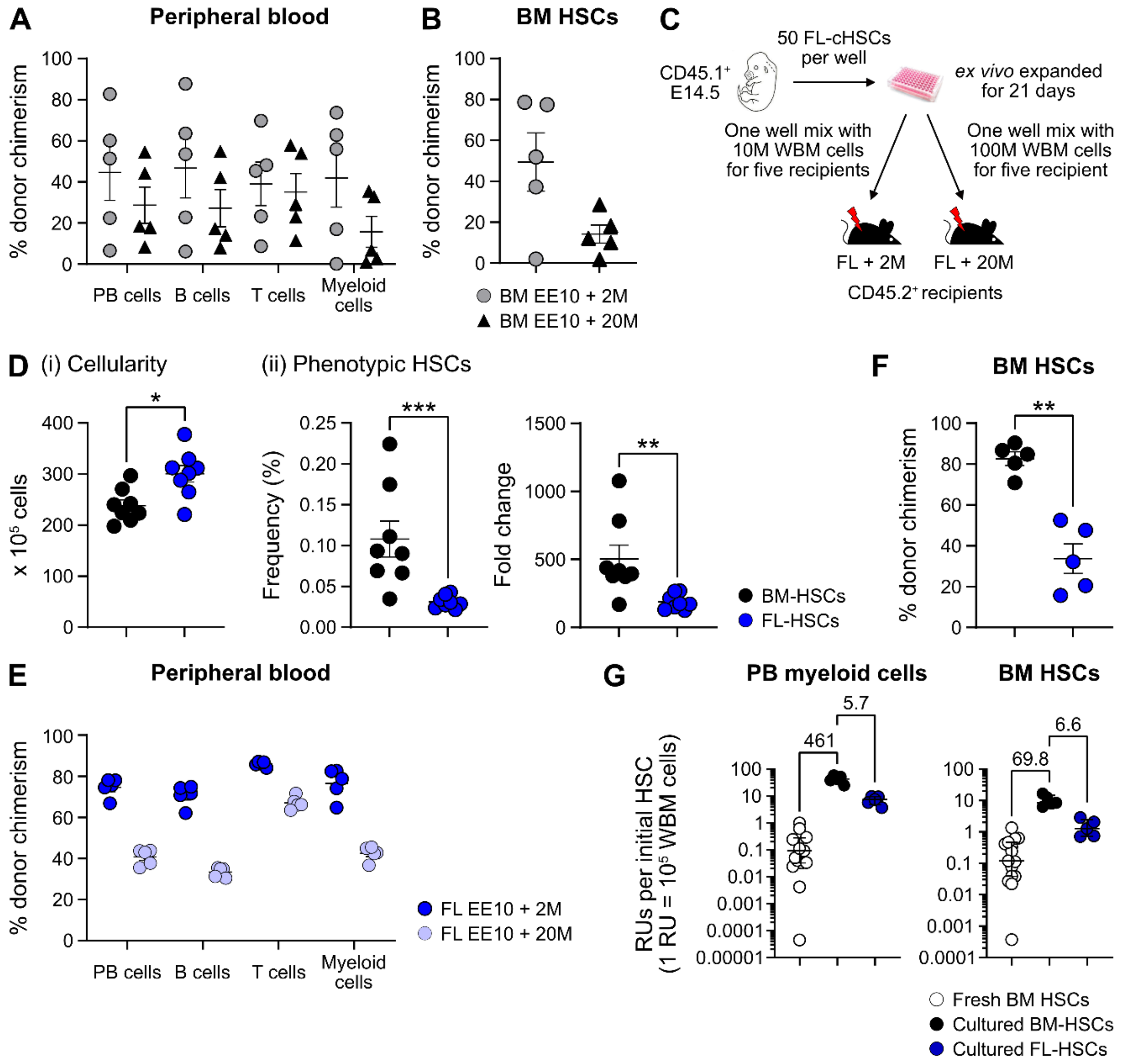


**Supplemental Figure 4. Quantification of HSC activity from cultured BM or FL HSCs**

(A) Percentage of test-cell derived cells in PB to each of the assessed lineages 16 weeks post-transplantation. Mice were transplanted with CD45.2$^+$ EE10 BM-cHSCs together with 2 or 20 million CD45.1$^+$ WBM cells.

(B) Test-cell derived chimerism in BM cHSCs 16 weeks post transplantation. Mice were transplanted with CD45.2$^+$ EE10 BM-cHSCs together with 2 or 20 million CD45.1$^+$ WBM cells.

(C) Outline of the competitive transplantation strategy to assess the repopulation ability of *ex vivo* expanded FL-cHSCs.

(D) Phenotypic analysis of *ex vivo* expanded BM-HSCs and FL-HSCs. (i). Overall cell expansion from 50 BM or FL $EPCR^{high}$ SLAM LSKs after 21-days of *ex vivo* culture. (ii). Frequency and fold change of phenotypic cHSCs ($EPCR^{high}$ SLAM LSKs) in *ex vivo* cultures after 21 days of culture from BM or FL-cHSCs.

(E) Test-cell derived chimerism in PB myeloid cells 16 weeks post transplantation.

(F) Test-derived HSCs chimerism in the BM of the recipients received *ex vivo* expanded cells from BM or FL-cHSCs 16 weeks post transplantation.

(G) RUs equivalent to one initial cHSC within PB myeloid cells and BM cHSCs in the recipients of 50 BM cHSCs or *ex vivo* expanded cells from 10 BM or FL-cHSCs 16 weeks post transplantation. Fold changes of EE10 BM-cHSCs versus 50 fresh BM cHSCs or EE10 FL-cHSCs are indicated. Median values are shown with interquartile ranges.

All data points depict values in individual recipients or culture wells. Error bars denote SEM. The asterisks indicate significant differences. *, $p < 0.05$; **, $p < 0.01$; ***, $p < 0.001$.

### Supplemental Figure 5

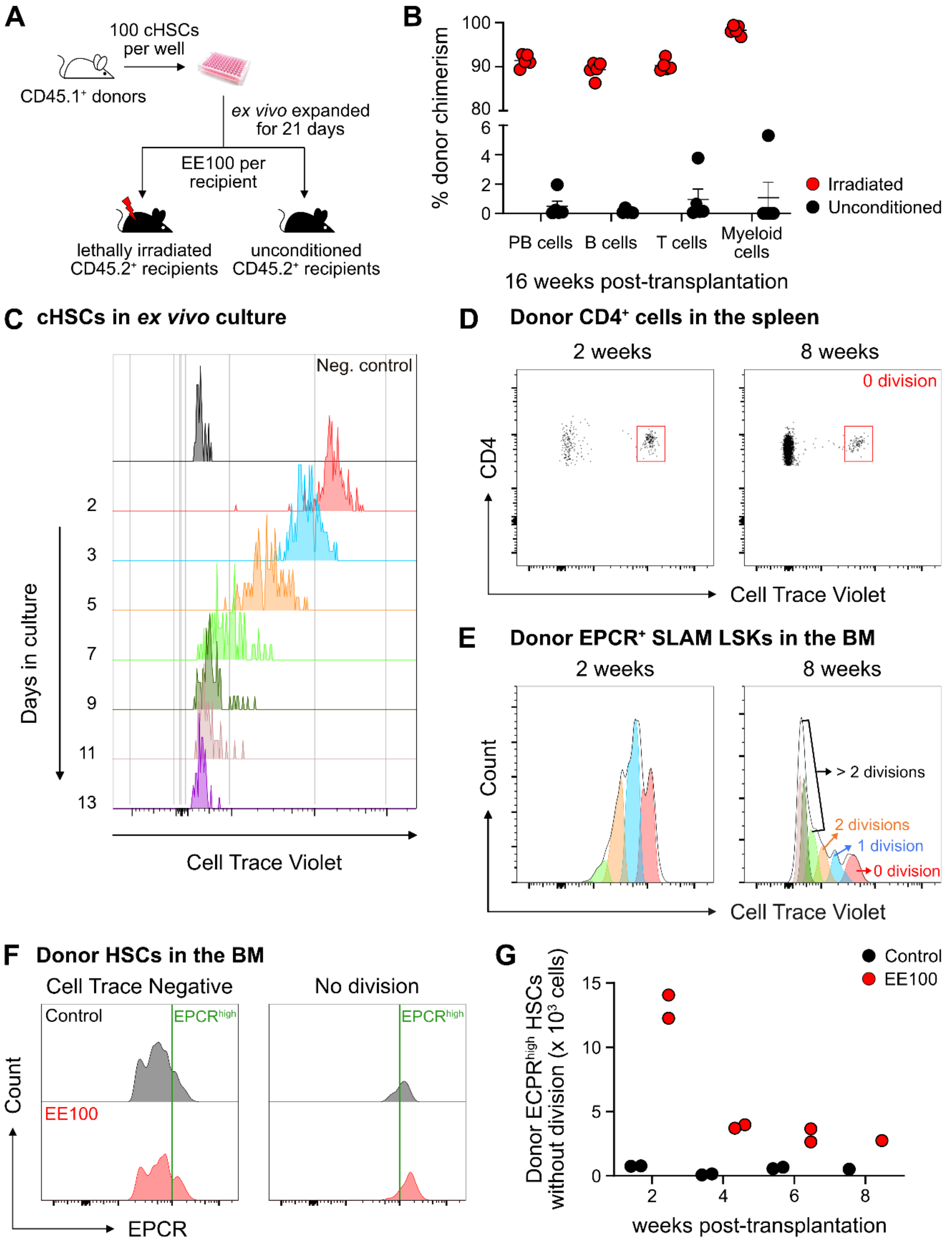


**Supplemental Figure 5. The fate of *ex vivo* cultured cHSCs following transplantation into unconditioned hosts.**

(A) Strategy to assess the repopulation of *ex vivo* expanded cHSCs into lethally irradiated or unconditioned recipients.

(B) Test-cell derived PB reconstitution 16 weeks post-transplantation. All data points depict values in individual recipients. Error bars denote SEM.

(C) CTV signal from cHSCs in ex vivo cultures. The cultures were initiated with 100,000 CTV labelled cKit-enriched cells per well (n = 5). Unstained cKit-enriched cultures were used as negative control. CTV signal was traced by analyzing half of the expanded cells after each split until 13 days after culture.

(D) CTV signal from transplanted CD4+ spleen cells, used to define undivided cells.

(E) Investigation of cell divisions based on CTV signals.

(F) Correlation of EPCR expression levels and the proliferative activity of cHSCs.

(G) Number of undivided donor EPCRhigh HSCs. All data points depict values in individual recipients.
